# Converging evidence of positive selection at height-associated loci in Europe

**DOI:** 10.64898/2026.01.27.702172

**Authors:** Valentin Hivert, Pierrick Wainschtein, Matthew R. Robinson, Evan K. Irving-Pease, Eske Willerslev, Michael E. Goddard, William Barrie, Julia Sidorenko, Peter M. Visscher, Loic Yengo

## Abstract

The contribution of natural selection to height differences across Europe remains elusive. Prior evidence of selection on height based on genome-wide association study (GWAS) data has been challenged because of residual confounding in earlier GWAS results. Here, we reassess this evidence using complementary approaches integrating multi-ancestry and within-family GWAS data with allele frequency estimates from 13 European ancestry populations. We detect signatures of positive selection at height-associated loci within and across populations, which remain significant after accounting for ancient admixture events and is largely driven by allele frequency differences between the Netherlands and other European ancestry populations. Altogether, our results provide new and converging evidence of polygenic adaptation on height in Europe.

## Introduction

Height exhibits a latitudinal gradient across Europe, with individuals from northern areas being, on average, taller than those from southern regions^1^. This observation largely aligns with Bergmann’s rule,^2^ stating that larger body sizes are expected in colder climates and has led to longstanding hypotheses regarding the role of natural selection in shaping height variation. Height is a highly heritable trait (with an estimated heritability of ∼80%)^3^ and is genetically correlated with the risk of several diseases^4^ such as cardiovascular disease^5,6^ and type 2 diabetes^7^, making its evolutionary trajectory of interest for both human genetics (understanding human evolution and genetic variation) and precision medicine.

Between 2012 and 2018, multiple studies have used summary statistics of genome-wide association study (GWAS) ^8–13^ of height and reported genetic evidence of polygenic adaptation for height in European ancestry (EUR) populations. However, subsequent analyses have also demonstrated that these findings were confounded by uncorrected population stratification,^14,15^ thereby weakening the conclusions of these previous studies. For example, when GWAS summary statistics from the UK Biobank (UKB) were used, evidence of selection either disappeared or was significantly attenuated^14,15^. In this context, Chen & Chiang^16^ provided important insights by showing that the issue of population stratification within EUR GWAS meta-analyses such as that conducted by Wood et al.^17^ extends across continents and genetic ancestries. Nevertheless, by combining data from the UKB and Biobank Japan and methods based on genetic differentiation (*F*_ST_), they detected selection signals at height-associated loci across continental populations, thus replicating previous results from Guo and colleagues^10^. These findings imply that natural selection may have acted on standing genetic variation globally, even if local signals remain difficult to disentangle from demographic history.

Since then, new GWAS resources based on larger sample sizes have become available. To date, Yengo et al.^18^ performed the largest and most genetically diverse GWAS of human height in over 5 million individuals. They identified 12,111 independent SNPs explaining approximately 40% of height variance in EUR populations. In addition, large within-family GWAS analyses, which are more robust to population stratification, have also become available and have been used to detect recent selection on height in ancestors of contemporary UK populations^19^. Additionally, a recent study leveraging ancient DNA data has suggested historical admixture rather than ongoing selection as a plausible cause of allele frequency differences observed at height-associated loci between modern EUR populations, although recent adaptive pressures cannot be ruled out^20^.

In this study, we use complementary approaches to reassess the potential role of selection in shaping height variation within Europe by leveraging summary statistics from the largest available GWASs of height^18,19^. We detect signals of recent selection at height-associated SNPs using trait-specific Singleton Density Score (tSDS) analyses^19^ and provide new evidence of local adaptation across 13 European populations using the 𝑄_X_ statistic^8^.

## Results

### Evidence of polygenic adaptation in ancestors of contemporary UK populations

The singleton density score (SDS) method was previously introduced to detect changes in allele frequencies within a population due to natural selection over recent evolutionary times (typically, within the past 2000 to 3000 years).^9^ An extension of the method, namely the trait-specific SDS (tSDS) method, combines GWAS summary statistics and SDS values at trait-increasing alleles to detect recent events of natural selection acting on trait-associated variants. Importantly, the tSDS method only uses the sign of the association and not the magnitude of the effect detected in GWAS. Using summary statistics from within-family GWAS and precomputed SDS in the UK population^9^, Howe et al.^19^ reported a positive correlation between the marginal GWAS p-values and tSDS, which provides evidence of positive selection for height-increasing alleles in ancestors of contemporary UK populations. We replicated their result using a slightly different approach focusing on testing if the mean tSDS of height-associated SNPs is significantly different from 0, as done previously. ^9,14,15^ A significantly positive mean tSDS implies positive selection, while a significantly negative mean tSDS would be expected under purifying selection or selection for the trait-decreasing alleles.

We first identified 228 independent SNPs (p-value < 5 × 10^-6^) from the Howe et al.^19^ EUR GWAS summary data using conditional and joint association analysis (COJO)^21^ and computed their tSDS values (using precomputed SDS in the UK population^9^). We found a marginally significant positive mean tSDS across these 228 SNPs (see Figure 1-A, mean tSDS = 0.013, p = 0.032), consistent with a signal of directional selection for taller individuals. We repeated this analysis using 7,630 independent genome-wide significant SNPs associated with height in a large GWAS meta-analysis of EUR cohorts,^18^ and found, consistently, a significantly positive mean tSDS across these SNPs (see Figure 1-B, tSDS mean = 0.084, p-value = 1.4 × 10^-13^). In addition to using summary statistics from within-family GWAS, we further sought to minimise confounding due to population stratification within Europe by replicating our findings using summary statistics from GWAS conducted in other ancestry groups.^18^ First, we demonstrated an extremely low level of residual EUR population stratification across our various sets of GWAS summary statistics, with the non-EUR and within-family GWAS exhibiting the lowest levels of such a residual stratification (see Supplementary Note 1). Using 358 and 1,187 height-associated SNPs identified from GWAS conducted in African ancestry (AFR) and Hispanic ethnicity (HIS), respectively, we also found significant evidence of positive selection on height (see Figure 1-C and D; tSDS mean = 0.202, p-value = 3.3 × 10⁻^4^ for AFR, and tSDS mean = 0.097, p-value = 1.4 × 10⁻³ for HIS). However, we could not replicate these results using 685 height-associated variants from a GWAS conducted in East Asian ancestry (EAS) populations, despite a positive mean tSDS value (see Figure 1-E, tSDS mean = 0.060, p-value = 0.14). Overall, our results are concordant with prior SDS-based studies conducted in the UKB^14,15^ and provide stronger statistical evidence of recent selection for height-increasing alleles in the UK by leveraging summary statistics from more powerful GWASs.

**Figure 1:**
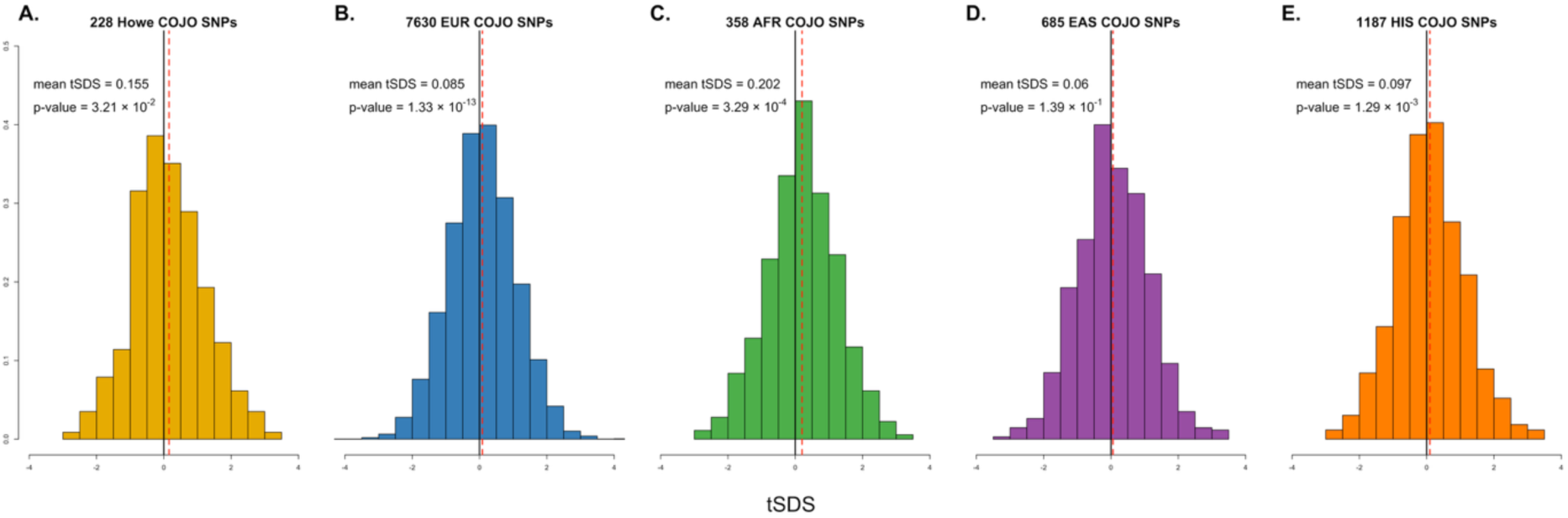
tSDS distributions across five independent sets of height-associated COJO SNPs. tSDS distributions are shown for (A) 228 EUR COJO SNPs from Howe et al.^19^ (within-family GWAS), as well as (B) 7,630 EUR, (C) 358 AFR, (D) 685 EAS and (E) 1,187 HIS COJO SNPs from Yengo et al.^18^. For each set of COJO SNPs, the mean tSDS value is depicted by the vertical red dashed line. P-values for one-sample t-test testing for departure from zero (vertical solid black line) are also indicated.

### Evidence of polygenic adaptation across Europe

The tSDS analysis is specifically designed to detect signals of polygenic adaptation occurring in the history of a single population. Therefore, we used an independent approach leveraging genetic information to detect signals of polygenic adaptation across multiple populations. We used the 𝑄_X_ statistics^8^ (see Material and Methods), an extension of the Lewontin and Krakauer test^22^ for polygenic scores, quantifying the over-dispersion of estimated genetic mean differences compared to a null model of genetic drift. We used allele frequency data at approximately one million SNPs from 13 EUR populations (see Material and Methods and Supplementary Table 1) along the European latitudinal gradient (average pairwise 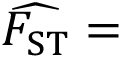 0.002, see Supplementary Figure 2) and estimated the mean polygenic score for height in each population using 7,172 COJO SNPs detected in a EUR GWAS (see Material and Methods). For a given set of summary statistics, the 𝑄_X_ p-value was calculated using a null distribution generated from 10,000 simulations, thus making 10⁻⁴ the minimum detectable p-value (see Materials and Methods).

Overall, we found a strong signal of polygenic adaptation across Europe (see Figure 2-A, 𝑄_X_= 318.21, p-value < 10^-4^), which was not driven by a single population as confirmed by our leave-one-population-out analyses (see Figure 2-B and Table 1). However, the Netherlands appeared to be an outlier as removing allele frequencies estimated in Dutch cohorts from the analysis halved the test statistic (𝑄_X_ = 166.42, p-value < 10^-4^).

**Figure 2:**
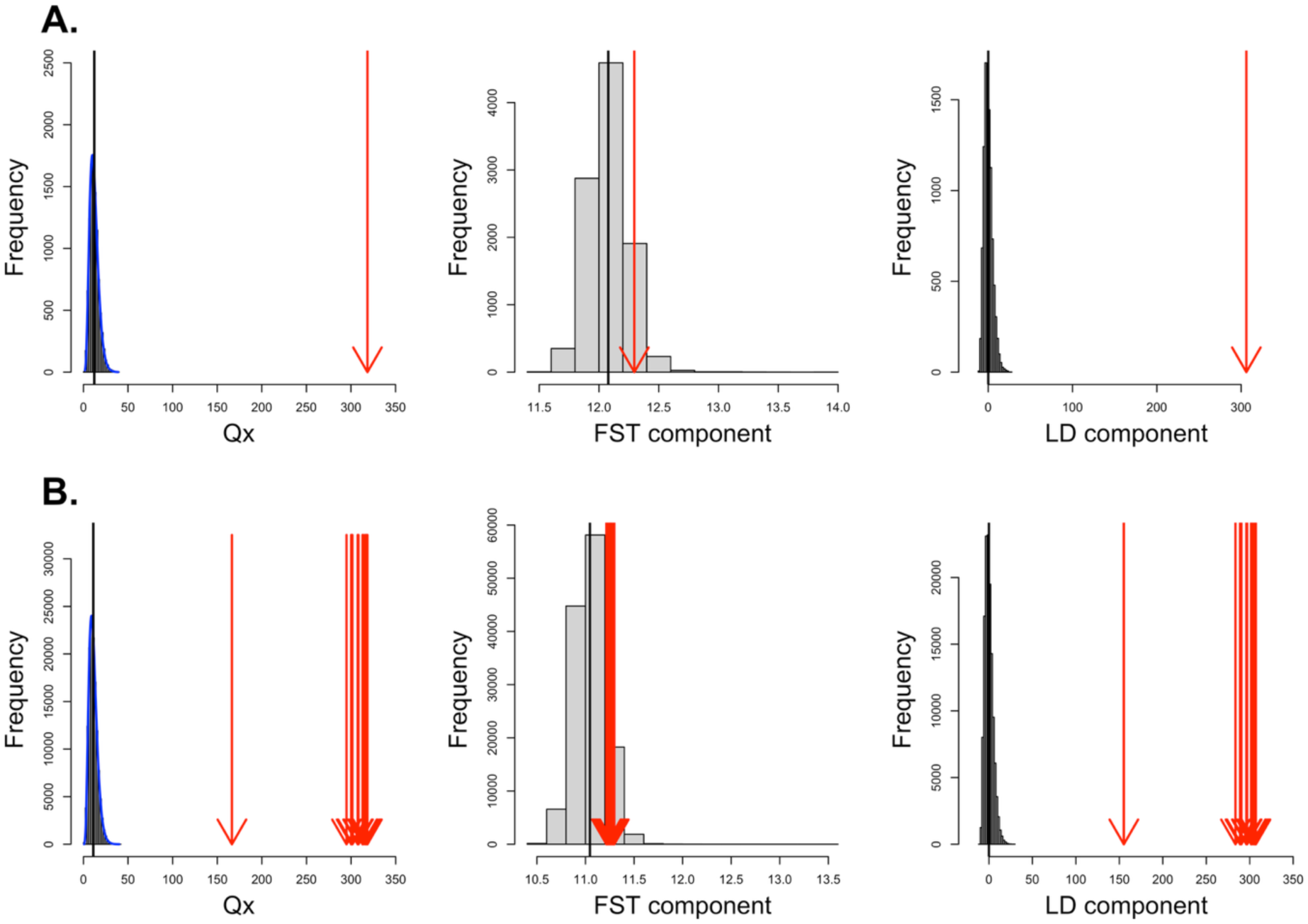
𝑸_𝐗_ **analysis of 7,172 SNPs associated with height in European ancestry GWAS.** Histogram of the empirical null distribution of 𝑄_X_, as well as its *F*_ST_ and LD component are displayed for A) analysis including the entire set of 13 European populations and B) leave-one-population-out analysis. Null distributions were obtained by genome-wide resampling of SNPs matched to the COJO SNPs by MAF and LD-score. Estimated values are depicted with red arrows and estimated mean values of the null distributions by the vertical black lines. The theoretical density of 𝑄_X_(see Material and Methods) is shown as a blue curve.

**Table 1:** 𝑸_𝐗_ **statistics (p-values) for analysis including 13 European populations and leave-one-population-out analysis across five independent set of height associated COJO SNPs.** For each set of populations, the 𝑄_X_ value and associated p-value (when p-value ≥ 10^-4^) is shown for European (EUR), African (AFR), East Asian (EAS) and Hispanic (HIS) COJO SNPs from Yengo et al.^18^, as well as EUR COJO SNPs from Howe et al.^19^ within-family GWAS.

| Population removed | SNP set<br>(Number of SNPs) |  |  |  |  |
| --- | --- | --- | --- | --- | --- |
|  | Yengo EUR<br>(7,172 SNPs) | Yengo AFR<br>(371 SNPs) | Yengo EAS<br>(676 SNPs) | Yengo HIS<br>(1,100 SNPs) | Howe EUR<br>(228 SNPs) |
| None (13 Populations) | 318.51 | 46.59 | 43.00 | 73.11 | 49.80 |
| Portugal | 294.86 | 44.16 | 31.62<br>($8 \times 10^{-4}$) | 68.10 | 45.89 |
| Spain | 301.53 | 46.72 | 42.03 | 68.58 | 48.79 |
| Italy | 300.10 | 46.40<br>($10^{-4}$) | 40.05<br>($10^{-4}$) | 59.04 | 46.93 |
| France | 318.34 | 41.10 | 42.31 | 66.77 | 46.80 |
| Switzerland | 308.23 | 44.66 | 40.78 | 70.11 | 44.22 |
| Great Britain | 301.20 | 42.65 | 40.73<br>($10^{-4}$) | 64.95 | 42.81 |
| CEU | 312.90 | 46.49 | 40.73 | 71.53 | 48.05 |
| Finland | 307.36 | 45.69 | 40.39 | 70.79 | 47.10 |
| Netherlands | 166.42 | 33.99<br>($3 \times 10^{-4}$) | 30.37<br>($1.2 \times 10^{-3}$) | 51.72 | 39.00 |
| Denmark | 312.93 | 46.48 | 42.62<br>($10^{-4}$) | 72.62 | 48.92 |
| Norway | 313.65 | 45.54 | 42.55 | 72.05 | 48.84 |
| Estonia | 317.11 | 46.41 | 43.02 | 67.72 | 48.46 |
| Sweden | 315.65 | 46.62 | 42.98 | 72.37 | 47.48 |

Consistently, partitioning the 𝑄_X_ statistic into marginal population-specific contributions (see Material and methods) confirmed the Netherlands as main outlier, contributing 87% of the 𝑄_X_signal of local adaptation across the 13 EUR populations (using Yengo et al.^18^ EUR COJO SNPs). The second largest contribution to the 𝑄_X_ statistic across all 13 populations was 14% from Portugal. Importantly, we did not find a significant linear association between mean polygenic score and latitude, consistent with analysis using summary statistics from previous GWAS meta-analyses^8^.

We replicated our findings using height-associated SNPs ascertained from Howe et al.^19^ (within-family GWAS) as well as GWAS conducted in non-EUR cohorts (see Table 1 and Supplementary Figures 3 to 6). These analyses also supported evidence of natural selection, albeit with reduced signals, consistent with the lower number of SNPs included and reduced predictive ability of these sets of SNPs^18^. The 𝑄_X_ statistics can be partitioned into two distinct components. The first, analogous to *F*_ST,_ captures non-directional genetic differentiation between populations at the GWAS SNPs. The second captures directional covariance of between-population allele frequency differences at GWAS SNP weighted by their effect sizes

(the LD-like component, see Material and Methods). Overall, we found the 𝑄_X_ signal to be mainly driven by the LD-like component, however, it is interesting to note that the analysis based on Howe et al. COJO SNPs were the only ones to display a relatively strong *F*_ST_-like signal between Sweden and the rest of Europe (the *F*_ST_-like component becomes not statistically significant when excluding Sweden from the analysis, see Supplementary Figure 6).

### Effect of ancient admixture

Irving-Pease et al.^20^ recently suggested that preferential migration of genetically-differentiated ancient Eurasian lineages towards Southern Europe (Farmer ancestry) versus Northern Europe (Steppe ancestry) contributes to the gradient of phenotypic and genetic mean differences observed between modern European populations. Importantly, the causes of the genetic differentiation between those ancient lineages remain largely unknown and could be explained by older selection events.

We sought to investigate the effect of differential admixture on our results by extending the 𝑄_X_ framework to include quantitative covariates (see Material and methods). We used this extension to decompose 𝑄_X_into contributions from allele frequency differences correlated with admixture proportions of ancient Eurasian lineages^20^ and a residual contribution independent of admixture. Across the range of GWAS summary statistics used in our analyses, we found that 43 to 87% of the 𝑄_X_ statistic was explained by differential admixture (see Supplementary Table 2). However, the overdispersion in allele frequency independent of admixture remained highly significant in most analysis (see Table 2), suggesting continued selection beyond the known differentiation between these ancient lineages. Similar to the previous analysis, we computed the marginal population-specific contributions to the 𝑄_X_ statistic after adjusting for differential admixture (see Figure 3). This again identified the Netherlands as the main outlier (72%), now followed by Great Britain (12%), in contrast to Portugal before adjustment.

**Figure 3:**
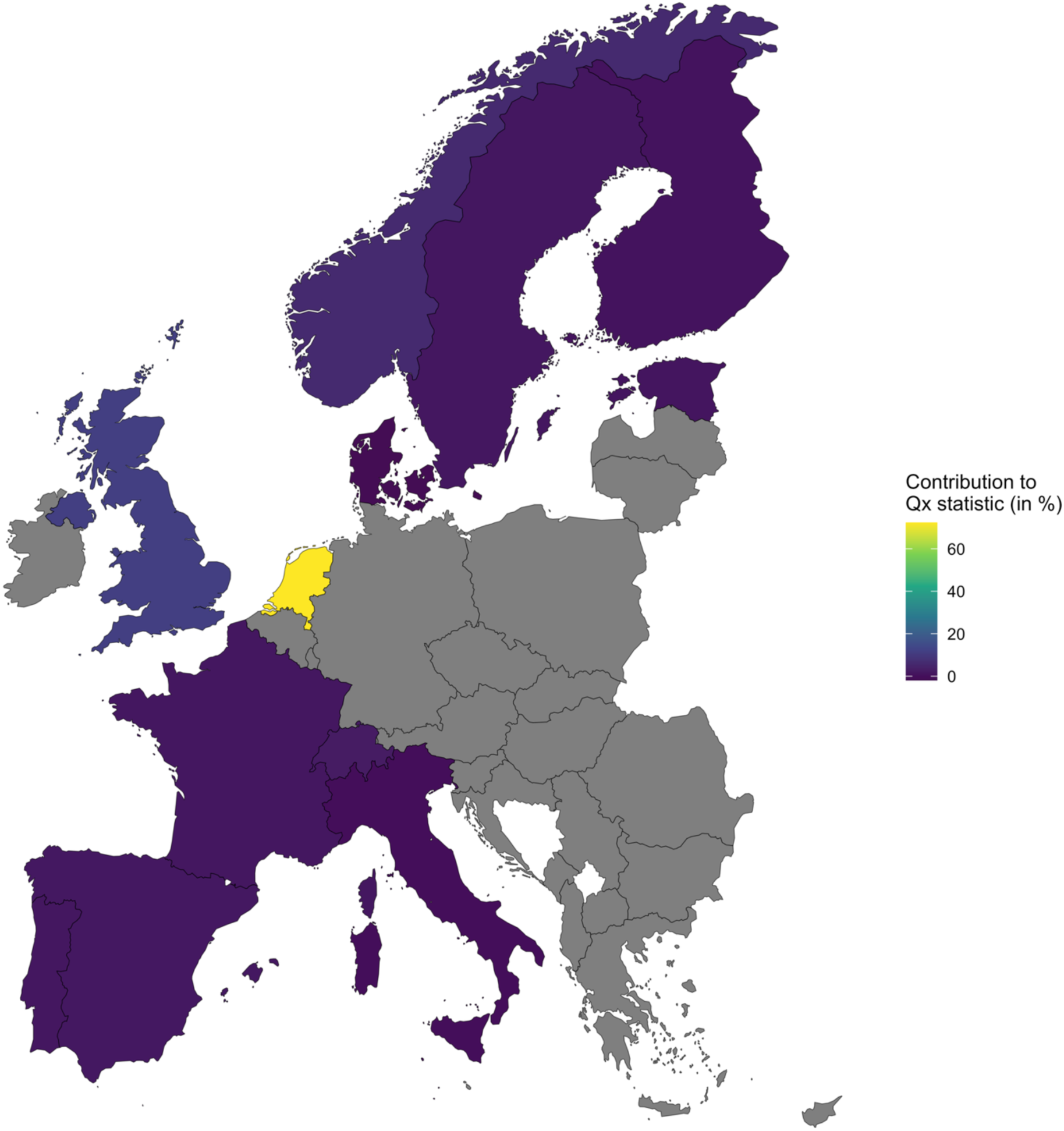
Population-specific contribution to the **𝑸_𝐗_**statistics including 6 admixture PCs as covariates and using European (EUR) COJO SNPs from Yengo et al.^18^. **For each** population, the marginal contribution to the 𝑄_X_ statistics (see Material and Methods) is shown in percentage.

**Table 2:** 𝑸_𝐗_ **statistics (p-values) including 6 admixture PCs as covariates for analysis including 12 European populations and leave-one-population-out analysis across five independent set of height associated COJO SNPs.** For each set of populations, the 𝑄_X_value and associated p-value (when p-value ≥ 10^-4^) is shown for European (EUR), African (AFR), East Asian (EAS) and Hispanic (HIS) COJO SNPs from Yengo et al.^18^, as well as EUR COJO SNPs from Howe et al.^19^ within-family GWAS.

| Population removed | SNP set<br>(Number of SNPs) |  |  |  |  |
| --- | --- | --- | --- | --- | --- |
|  | Yengo EUR<br>(7,172 SNPs) | Yengo AFR<br>(371 SNPs) | Yengo EAS<br>(676 SNPs) | Yengo HIS<br>(1,100 SNPs) | Howe EUR<br>(228 SNPs) |
| None (12 Populations) | 132.97 | 17.41<br>($3.9 \times 10^{-3}$) | 20.99<br>($6 \times 10^{-4}$) | 22.21<br>($7 \times 10^{-4}$) | 27.71<br>($1 \times 10^{-4}$) |
| Portugal | 123.56 | 12.33<br>($1.6 \times 10^{-2}$) | 11.43<br>($2.2 \times 10^{-2}$) | 19.09<br>($8 \times 10^{-4}$) | 20.03<br>($4 \times 10^{-4}$) |
| Spain | 123.20 | 11.35<br>($2.1 \times 10^{-2}$) | 11.31<br>($2.3 \times 10^{-2}$) | 18.41<br>($1.2 \times 10^{-3}$) | 19.68<br>($8 \times 10^{-4}$) |
| Italy | 133.37 | 16.59<br>($2.9 \times 10^{-3}$) | 17.54<br>($1.4 \times 10^{-3}$) | 22.33 | 23.97 |
| France | 131.10 | 16.16<br>($1.8 \times 10^{-3}$) | 20.86<br>($3 \times 10^{-4}$) | 21.63 | 27.45 |
| Switzerland | 124.61 | 16.20<br>($2.6 \times 10^{-3}$) | 15.26<br>($2.9 \times 10^{-3}$) | 21.65 | 23.64 |
| Great Britain | 106.74 | 13.33<br>($1.1 \times 10^{-2}$) | 18.97<br>($6 \times 10^{-4}$) | 16.48<br>($2.2 \times 10^{-3}$) | 20.31<br>($3 \times 10^{-4}$) |
| Finland | 88.81 | 14.41<br>($4.8 \times 10^{-3}$) | 16.06<br>($2.7 \times 10^{-4}$) | 17.17<br>($1.5 \times 10^{-3}$) | 27.27 |
| Netherlands | 25.12<br>($1 \times 10^{-4}$) | 8.15<br>($8.2 \times 10^{-2}$) | 10.51<br>($3.1 \times 10^{-2}$) | 7.19<br>(0.12) | 20.31<br>($8 \times 10^{-4}$) |
| Denmark | 79.00 | 12.08<br>($1.8 \times 10^{-2}$) | 17.10<br>($1.3 \times 10^{-3}$) | 14.04<br>($6.7 \times 10^{-3}$) | 26.99 |
| Norway | 99.94 | 8.34<br>($7.8 \times 10^{-2}$) | 11.79<br>($1.9 \times 10^{-2}$) | 12.22<br>($1.6 \times 10^{-2}$) | 6.12<br>(0.19) |
| Estonia | 107.09 | 16.63<br>($2.3 \times 10^{-3}$) | 18.31<br>($10^{-3}$) | 19.73<br>($7 \times 10^{-4}$) | 26.58<br>($10^{-4}$) |
| Sweden | 127.37 | 16.96<br>($1.8 \times 10^{-3}$) | 19.32<br>($4 \times 10^{-4}$) | 21.11<br>($2 \times 10^{-4}$) | 21.19<br>($6 \times 10^{-4}$) |

To further explore the timing and strength of selection, we used PALM^23^ (Polygenic Adaptation Likelihood Method), an ancestral recombination graph (ARG) based approach designed to infer selection over specific time windows. While the original PALM study found no significant selection on height over the last 2,000 years using UK Biobank and GBR individuals, we extended this analysis using COJO summary statistics from European (EUR), African (AFR), and East Asian (EAS) populations. Overall, we observed a marginally significant selection gradient (𝜔 = 0.020 ± 0.0095, p = 0.04) translating into an average per-SNP selection coefficient for trait increasing allele 𝑠 = 2.16 × 10^-4^ over the last 2.7k years (or approximately 93 generations assuming 29 years per generation) using the EUR GWAS (see Supplementary Table 3) but not with other GWAS datasets.

## Discussion

Our findings provide converging lines of evidence supporting positive selection of height-increasing alleles in EUR populations. Using trait-specific singleton density score (tSDS) analyses, we detected significant signals of positive selection favouring height-increasing alleles in ancestors of contemporary populations in the United Kingdom. Our tSDS-based analyses corroborate prior observations that the frequency of alleles associated with increased stature have risen over the past 2,000 to 3,000 years in Northern Europe, reinforcing the hypothesis of recent directional selection on height^9^.

Further, 𝑄_X_-based polygenic adaptation tests reveal robust local adaptation signals at height-associated loci across EUR populations. Importantly, these signals persist after accounting for population stratification and differential admixture with ancient Eurasian lineages, although they weaken. Nevertheless, ancient admixture explains a large proportion of the apparent signal of polygenic adaptation, consistent with both differential admixture and natural selection acting as drivers of the observed mean genetic differences. Interestingly, the Netherlands was an outlier in all our 𝑄_X_analyses, exhibiting elevated frequencies of height-increasing alleles relative to all other EUR populations. This result aligns with known phenotypic extremes, i.e., the Dutch being among the tallest populations globally^24^. However, earlier work has shown that the increase in mean height in the Dutch population over the last centuries cannot be explained by natural selection alone^25^. While selection has shaped height-related loci, additional factors such as assortative mating, environmental influences, and socioeconomic confounders likely interact with genetic architectures to produce the extreme height phenotype observed in the Dutch population.

We employed within-family GWAS designs and data from other ancestry groups to address prior concerns regarding confounding due to stratification in standard GWAS^14,15^. These approaches validate the presence of selection signals independent of stratification, reinforcing confidence in our conclusions. Berg et al.^14^ raised concerns about the validity of 𝑄_X_ in the presence of residual LD between GWAS variants in the analysis, which can inflate signals and distort the null distribution. To address this, we selected 1,489 COJO SNPs, choosing the most significant variant from each independent LD block based on the Yengo 2022 EUR summary statistics. This sensitivity analysis still yielded a strong signal (𝑄_X_=112.91, p-value < 10^-4^, see Supplementary Figure 7), exceeding the expected value from a random set of COJO SNPs (see Supplementary Figure 8). Following prior recommendations^14^, we also generated a null 𝑄_X_ distribution for the original Yengo EUR dataset by randomly flipping the sign of effect sizes, thus preserving within-chromosome LD structures. The resulting distribution closely matched the theoretical 𝜒_M-1_^2^ alleviating concerns about misspecification of the null. Assortative mating has also been proposed as a factor that could distort the null distribution of 𝑄_X_. To test this, we simulated multiple populations undergoing assortative mating on trait with an equilibrium heritability of 0.8 (e.g., height) and a spousal correlation r=0.2^26^ (see Materials and Methods). Over 100 simulations replicates, we showed that assortative mating can indeed increase the mean 𝑄_X_ value (see Supplementary Figure 9, genomic inflation factor 𝜆_12_ = 1.13 and 1.19 for 10 and 100 generations respectively).

However, even after 100 generations of assortative mating across 13 subpopulations, the resulting 𝑄_X_distribution remained well below our observed signal. This suggests that assortative mating alone is unlikely to explain the detected selection signal at height-associated loci. Our triangulation across independent methodologies and GWAS well controlled for population stratification provides a robust case for polygenic selection shaping height variation across Europe.

Future work should also investigate the sex specificity of natural selection, especially given sexual selection hypotheses for male height^27,28^. Examining latitudinal and sex-stratified selection patterns using sex-differentiated GWAS may yield further insight.

Finally, our findings have broader implications because height is genetically correlated with numerous diseases (for example, cardiovascular disease^5,6^ and type 2 diabetes^7^) such that selection on height-associated loci may have influenced disease genetic architectures.

In summary, our results support a key role of natural selection in shaping height differences across Europe, while emphasising the importance of methodological rigour and evolutionary context when interpreting polygenic adaptation. As richer datasets from ancient and global populations become available, our ability to untangle the evolutionary and biomedical consequences of selection will continue to improve.

## Material and methods

### Population genetics dataset

We compiled allele frequency data from 13 European countries using multiple publicly available datasets. We utilised data from the 1000 Genomes Project Phase 3 (1KGP), selecting 503 individuals from five European subpopulations: Tuscans (TSI), Iberians (IBS), British (GBR), Finns (FIN), and Utah residents of Northern and Western European ancestry (CEU). Additional genotype data were obtained from the POPRES study through the database of Genotypes and Phenotypes (dbGaP)^29^. As in previous work^12^, we selected 1,472 individuals from Portugal, Spain, Italy, France, and Switzerland whose four grandparents were born in the same country as the individual. Genotypes were aligned to the positive strand, and the genomic build was lifted from hg18 to hg19 using the command-line version of LiftOver, provided by the UCSC Genome Browser project^30^.

A total of 447,161 autosomal SNPs with minor allele frequency (MAF) > 0.01 were retained. After merging with 1KGP data and flipping non-ambiguous alleles using conform-gt^31^, 377,584 SNPs remained. Genotype pre-phasing was performed using SHAPEIT5^32^, and imputation was carried out with BEAGLE5^33^ using the HRC reference panel (reference EGAD00001002729). Post-imputation quality control included removal of duplicate SNPs, poorly imputed SNPs with DR2 < 0.8, SNPs with missingness > 0.05, individuals with missingness > 0.1, SNPs with MAF < 0.01, and SNPs failing Hardy-Weinberg equilibrium (p-value < 1e-4). This yielded a final dataset of 6,253,830 SNPs shared with the 1KGP reference. We merged the IBS and TSI subpopulations from 1KGP with Spain and Italy respectively. We then used Country-Specific Allele Frequency sources:

- **Estonia**: Allele frequencies were obtained from 49,160 individuals of the Estonian Biobank (EEB)^34^.
- **Netherlands**: Frequencies were obtained from 64,439 European individuals of Lifelines: a population-based study to study disease overriding risk factors for the development of multifactorial diseases during lifetime, release 1 and 2 imputed on the HRC reference panel^35^.
- **Norway**: Frequencies were sourced from publicly available GWAS summary statistics based on the MoBa cohort^36^.
- **Denmark**: Data were obtained from publicly available GWAS summary statistics based on the iPSYCH cohort^37^.
- **Sweden**: Frequencies were taken from the SweGen project (release 20240627), generated by Science for Life Laboratory^38^.

Each population specific dataset was ascertained for the set of European variants present in the curated dataset made from the combination of POPRES and 1KGP. We retained a final list of 890,392 HapMap3 SNPs common to all datasets and for which pre-computed LD scores were available^39^. Allele frequency comparisons across datasets, including POPRES and 1KGP, are shown in Supplementary Figures 10 and 11, and sample sizes used to estimate those allele frequencies are shown in Supplementary Table 1.

### COJO summary statistics

We utilised ancestry-specific summary statistics derived from conditional and joint analysis (COJO)^21^ multiple SNPs analysis as implemented in the GCTA software^40^. Specifically, we used publicly available COJO summary statistics from Yengo et al.^18^, comprising 7,630 significant and approximately independent SNPs from individuals of European (EUR) ancestry with pre-computed SDS values, and 7,172 in the HapMap phase 3 panel (HM3) present in our population genetic SNP set. Additionally, we included 371 (358) SNPs from African (AFR), 676 (685) from East Asian (EAS), and 1,100 (1,187) from Hispanic (HIS) ancestry groups in 𝑄_X_ (tSDS) analysis, selected based on their genome-wide significance and approximate independence.

To complement these data, we also performed COJO analysis using publicly available summary statistics from the within-family GWAS conducted by Howe et al.^19^. To obtain a LD reference panel, we analysed a large dataset of 347,979 unrelated (genomic relatedness< 0.05) individuals of European ancestries from the UK Biobank (UKB) using the genetic ancestry calling from Wang and colleagues^41^. Informed consent was obtained from all study participants and those who expressed the wish to be withdrawn were removed from analysis. We used the release 3 of the UKB where individuals were genotyped on the Affymetrix UK Biobank Axiom array before imputation using the HRC and UK10K reference panel and IMPUTE2. SNPs were filtered for quality control in Europeans by removing those with missing genotyping rate > 0.05, Hardy-Weinberg equilibrium test p <1e-6, and MAF < 0.01. After filtering, we extracted autosomal HapMap phase 3 (HM3) markers, resulting in 1,173,448 SNPs. Finally, we used a random subset of 50K unrelated individuals as LD reference panel for COJO analysis, as well as a genomic window size of 10Mb, and a COJO threshold p-value of 5 × 10^-6^, identifying 228 significant and independent HM3 SNPs present in our population genetics dataset and for whom pre-computed SDS values were also available.

### Trait Singleton Density Score analysis

To investigate recent selection on height-associated loci, we performed a trait Singleton Density Score (tSDS) analysis using precomputed SDS values from UK10K, covering 4,441,773 SNPs^9^. For each set of COJO-derived GWAS summary statistics, we extracted the corresponding SDS values and polarised them with respect to the trait-increasing alleles to obtain tSDS values. We then tested whether the mean tSDS deviated significantly from zero using a one-sample t-test, with a positive mean indicating recent positive selection on height-increasing alleles.

### Detecting local adaptation with **𝑸_𝐗_** analysis

The 𝑄_X_ statistics is designed to test for polygenic adaptation across *M* populations using summary statistics. It compares observed variance in estimated genetic means among populations to the expectation under a neutral model of genetic drift, leveraging population allele frequency data and estimated SNP effect sizes from GWAS. For a quantitative trait with *L* independent causal loci, the genetic means in the population m is given by:

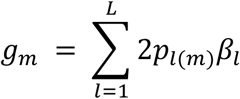

Where 𝑝_l(_*_m_*_)_ is the effect allele frequency of locus 𝑙 in population *m*, and 𝛽*_l_* its effect size. Under a null model of genetic drift, and assuming that causal variants and effects are shared across populations, the vector of population genetic means 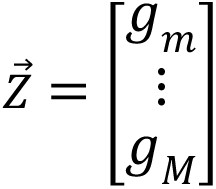 follow a multivariate normal distribution:

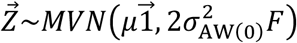

With 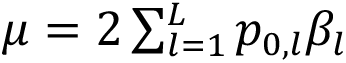 the ancestral population’s genetic mean, and 𝑝_o,l_ is the effect allele frequency of locus 𝑙 in the ancestral population (estimated as the average frequencies across populations). Additionally,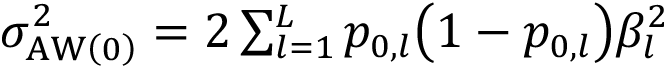 is the additive genetic variance within the ancestral population, and *F* the 𝑀 × 𝑀 matrix of variance-covariance of genome-wide allele frequencies. The 𝑄_X_ statistic is defined as:

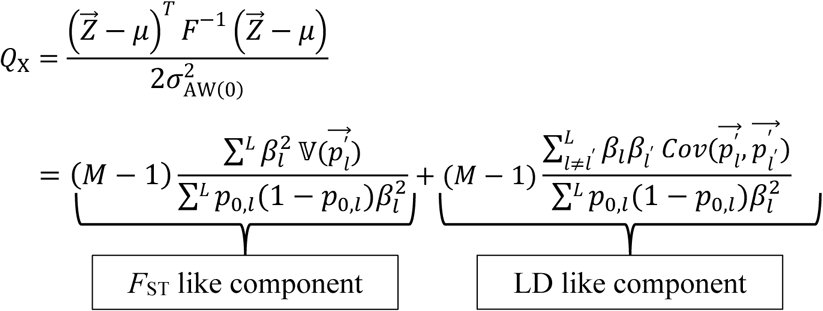

Where 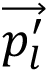 is a vector of transformed allele frequencies to control for population structure^8^. Under a null model of genetic drift, 𝑄_X_follow a 𝜒_M-1_^2^distribution. For a given set of summary statistics, we simulated the null distribution of 𝑄_X_(using 10,000 replicates) by randomly sampling putatively neutral variants genome-wide matched on allele frequency and LD-score^39^ with the GWAS SNPs.

We can further partition 𝑄_X_ into the sum of population-specific contributions:

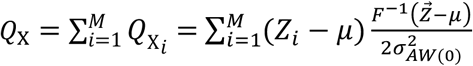

Where 𝑄_Xi_ represents the *F*_ST_ and LD-like components of 𝑄_X_ associated with population 𝑖. It therefore quantifies the marginal contribution of population 𝑖 to the overall statistic. The relative contribution of each population can be expressed as 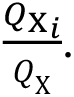 Importantly, these marginal contributions can be either positive or negative. A positive value indicates that population 𝑖 contributes to an excess of mean genetic differences relative to the expectation under the null model of genetic drift, whereas a negative value indicates that population 𝑖 reduces this departure.

As per previous recommendations to overcome the rank-deficiency of the matrix of variance-covariance *F* (see Berg et al.^8^ for details), we excluded one population (chosen at random) from the mean-centered vector of genetic means and matrix *F* when computing the 𝑄_X_statistic. However, this approach prevents us to compute the marginal contribution of each population. An equivalent alternative that retains all populations is to use the Moore–Penrose generalised inverse of the *F* matrix. In our analyses, we found both approaches to yield strictly equivalent results.

### Including admixture PCs in **𝑸_𝐗_** analysis

To enable the inclusion of quantitative covariates, we extended the 𝑄_X_ model to incorporate a design matrix *X* of *J* mean-centered population-level covariates. Assuming a linear model, we have:

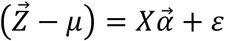

with 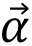 the vector of effect sizes and 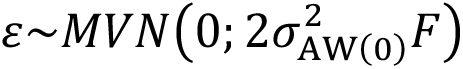 the residuals. The generalised least square solution is:

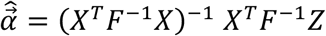

And residuals are estimated as:

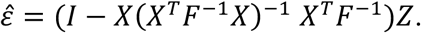

The covariate-adjusted 𝑄_X_ statistic becomes 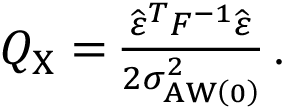 Under the null hypothesis of genetic drift, it now follows a 𝜒_M-j-1_^2^ distribution (see Supplementary Figure 12).

We used previously estimated admixture proportions of ancient Eurasian lineages in 12 European populations (excluding CEU)^20^. These lineages include Western Hunter-Gatherers (WHG), Eastern Hunter-Gatherers (EHG), Caucasus Hunter-Gatherers (CHG), Neolithic Farmers, Yamnaya Steppe, and ancient African and East Asian ancestries. We performed PCA on the raw admixture proportions and retained the first six PCs (capturing 100% of variation) as covariates in the 𝑄_X_analysis. The PCs were independently recalculated for every leave-one-population-out analysis.

### Polygenic Adaptation Likelihood Method (PALM) analysis in British ancestry

We investigated the timing and strength of selection at height-associated loci using PALM^23^, an ancestral-recombination-graph (ARG) based method that integrates population genetics data with GWAS summary statistics to infer positive selection across specified time windows. We inferred the coalescent trees with Relate^42,43^ v1.2.1 using Great Britain samples from 1KGP, assuming a mutation rate 𝜇 = 1.25 × 10^-8^ and a generation time of 29 years. We then applied PALM to test for evidence of selection on height-associated variants over the past 2,000 and 2,700 years, using Yengo et al.^18^ EUR, AFR and EAS COJO summary statistics.

### Simulation study with assortative mating

We simulated the evolution of a quantitative trait under assortative mating in *M* = 13 populations derived from a common ancestral population. In each population, we simulated *L* = 14,000 SNPs, half of which were causal for the trait driving assortment and half were independent of that trait. Ancestral allele frequencies were drawn from a uniform distribution (MAF ≥ 0.01). Effect sizes for causal SNPs were sampled from a normal distribution and scaled such that the additive genetic variance at equilibrium matched a target equilibrium heritability (ℎ*_eg_*^2^, that is after multiple generation of assortative mating) of 0.8. The phenotypic variance at equilibrium was 𝜎*_p,eq_*^2^ = 1. The additive genetic variance in the founding generation (𝜎*_A_*_(0)_^2^) was adjusted to account for the specified spousal correlation (*r*) following Eq. 7.19a of Lynch & Walsh^44^. Phenotypes were computed as the sum of additive genetic values and normally distributed environmental effects. Populations evolved for either 10 or 100 generations, with mating pairs formed according to phenotypic similarity to achieve the specified spousal correlation 𝑟 = 0.2 (see Stulp et al.^26^). Offspring inherited alleles under Mendelian segregation, and allele frequencies were recorded for all loci at the final generation. SNPs not associated with the trait driving assortment were utilised to estimate the matrix of variance-covariance of allele frequencies used in the 𝑄_X_ analysis. Simulation experiment was conducted using a custom R script with 1000 replicates per simulation setting. Inflation of the test statistic was quantified with the genomic inflation factor:

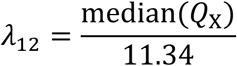

With median(𝑄_X_) denoting the median of 𝑄_X_ value across the 1000 replicates and 11.34 the expected median 𝜒_12_^2^, which is the expected distribution under the null.

## Data and code availability

Analysis were performed using R version 4.4.3^45^ and 𝑄_X_ analysis scripts were modified from the source code located at https://github.com/jjberg2/PolygenicAdaptationCode.

Scripts used for analysis are available on Github: https://github.com/vhivert/Natural-Selection-Height-Europeans

## Acknowledgments

L.Y. is supported by the Australian Research Council (FT220100069) and the Snow Medical Research Foundation. P.M.V. is supported by the Australian Research Council (FL180100072).

E.K.I.P is supported by the Royal Society (URF\R1\251675). This study makes use of data from the UK Biobank under the application number 12505, Lifelines under the application number OV19_0495, and dbgap POPRES data project #3189, accession phs000145.v4.p2. We wish to acknowledge The University of Queensland’s Research Computing Centre (RCC) for its support in this research. We thank John Novembre for valuable comments on an early version of the manuscript. We are also grateful to Bruce Walsh, Tian Lin, Clara Albinana, Tunde Olasege and Solal Chauquet for helpful discussions.

## Web resources

GCTA (v1.94.1), https://yanglab.westlake.edu.cn/software/gcta/#Overview RELATE: https://myersgroup.github.io/relate/index.html

PALM: https://github.com/standard-aaron/palm

Precomputed SDS from UK10K: https://datadryad.org/dataset/doi:10.5061/dryad.kd58f

