## Supplementary Material for "Converging evidence of positive selection at height-associated loci in Europe"

### Supplementary Note 1: Quantifying residual European population structure in human height GWASs

Residual population stratification (PS) is a major concern in GWAS, particularly following evidence of biased inference of natural selection on height in Europe from the Wood et al.[1] GWAS summary statistics. This bias was driven by uncorrected PS along the European latitudinal axis[2,3]. To evaluate whether residual PS may affect the GWAS datasets used in our study (we used the publicly available GWAS summary statistics that exclude 23andMe), we quantified it following the approach of Yengo et al.[4]. We computed the squared correlation between estimated marginal SNP effects and SNP loadings on the first 20 principal components (PCs) derived from 503 European samples from the 1000 Genomes Project (1KGP). The analysis was restricted to 101,360 independent HapMap3 SNPs, consistent with previous work[4]. For comparison, we included an extra set of European GWAS summary statistics, namely Wood et al.[1], Yengo et al.[5], but also UKB-456k (BOLT-LMM): a GWAS in 456,414 EUR participants of the UKB, UKB-350k (unrelated): a GWAS in 348,501 unrelated (i.e. estimated genetic relatedness <0.05) EUR participants of UKB, Within-family (UKB): a family-based GWAS performed in 17,942 independent EUR sibling pairs from the UKB, and Robinson et al.[6]: a family-based GWAS performed using 17,500 sibling pairs from Hemani et al.[7]. The analysis of the Robinson et al.[6] summary statistics included 70,316 SNPs out of the 101,360 independent HapMap3 SNPs.

As shown in Supplementary Figure 1, SNP loadings on PC2 explained the largest proportion of the variance in SNP effects across datasets, consistent with PC2 capturing the North–South genetic structure in Europe. However, the magnitude of this variance varied across GWASs. The Wood et al.[1] GWAS showed the highest stratification (~2.6% of variance explained by the first 20 PC loadings), while the other GWASs tested showed markedly lower levels (<0.5%). It is interesting to note that non-European GWAS showed levels of stratification similar to the within-family GWASs, except for the Hispanic GWAS that showed similar PS to the Yengo et al.[4] EUR GWAS. This can potentially be explained by high level of European admixture in the Hispanic cohort used here (that exclude 23andMe).

This result demonstrates that GWASs used in our study substantially mitigate stratification, with family-based and non-European GWASs providing the most robust control.

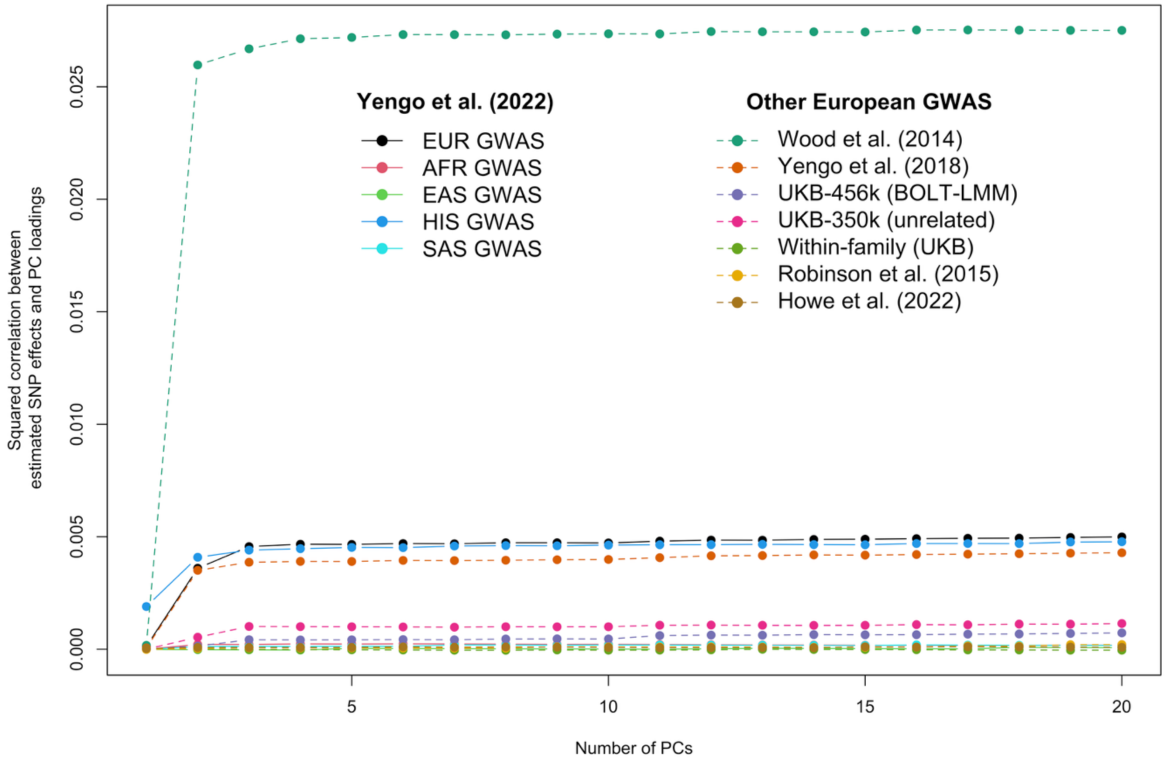


**Supplementary Figure 1: Quantification of residual European population stratification in different set of GWAS summary statistics for height.** We show the variance in estimated marginal SNP effects from various height GWAS that is explained by SNP loading on genotypic principal components (PC) calculated among 503 EUR samples from the 1KGP. The x-axis indicates the number of vectors of PC loadings included in the regression model (SNP effects regressed on PC loadings).

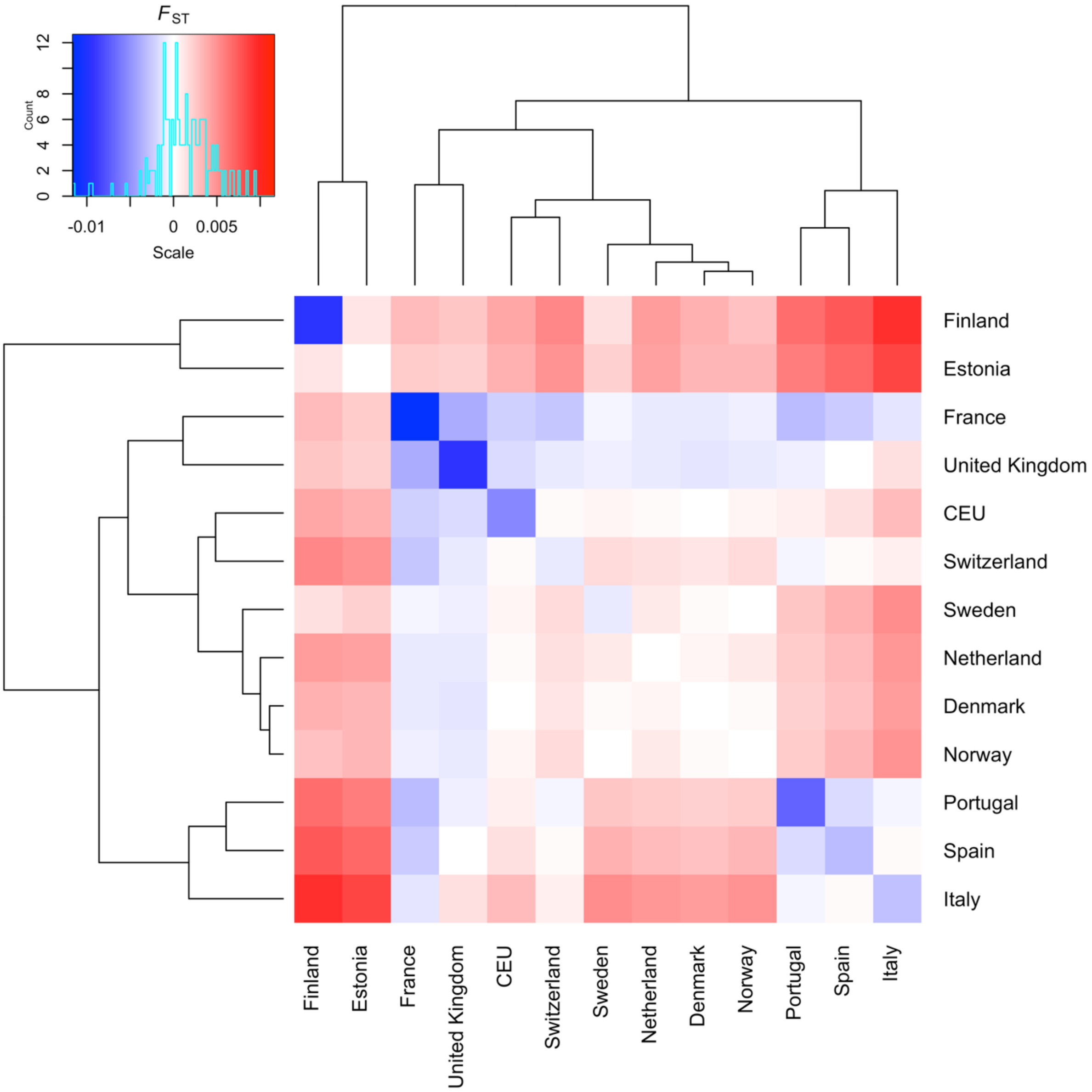


**Supplementary Figure 2: Estimated pairwise $F_{ST}$ across the 13 European countries included in the population genetics analysis.** Pairwise $F_{ST}$ were calculated from 50,000 randomly sampled HapMap3 SNPs using Hudson's estimator[8], as described in Bhatia et al.[9]

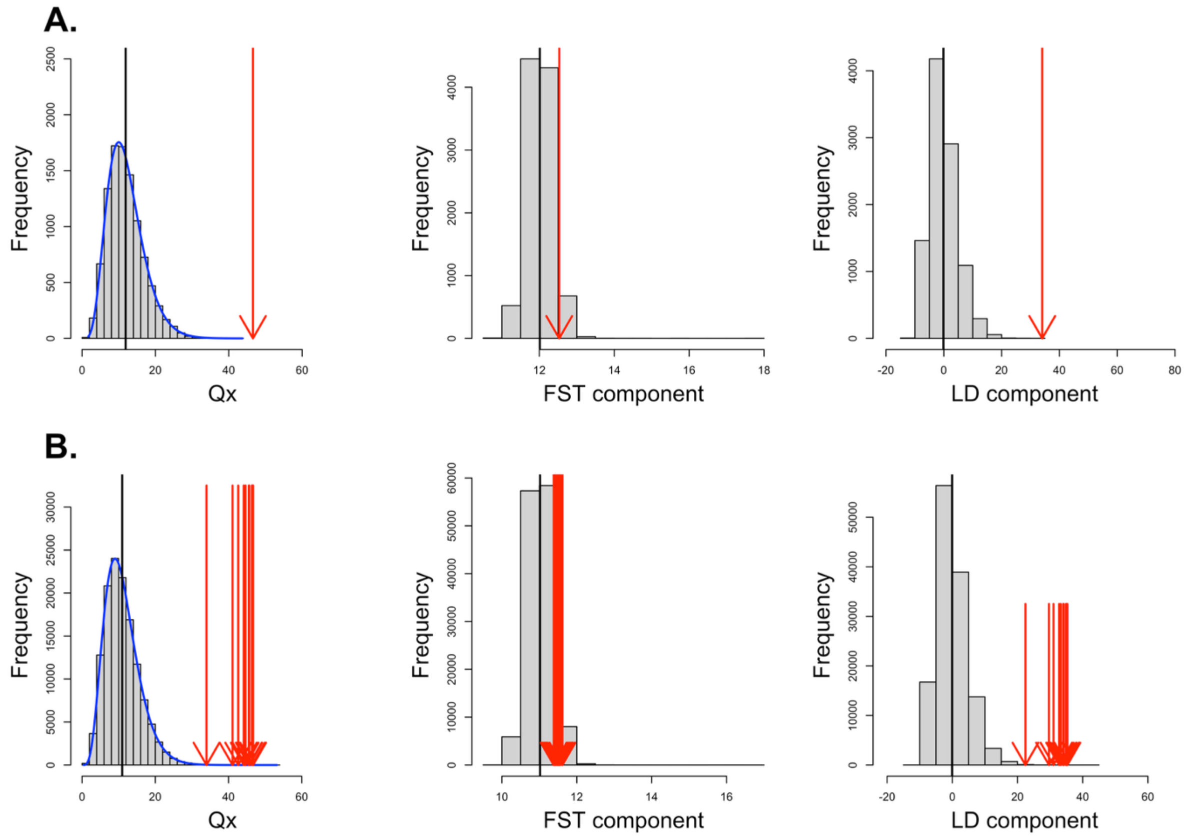


**Supplementary Figure 3: $Q_X$ analysis using 371 AFR COJO SNPs from Yengo et al.[4]**
Histogram of the empirical null distribution of $Q_X$, as well as its $F_{ST}$ and LD component are displayed for A) analysis including the entire set of 13 European populations and B) leave-one-population-out analysis. Null distributions were obtained by genome-wide resampling of SNPs matched to the COJO SNPs by MAF and LD-score. Estimated values are depicted with red arrows, and estimated mean values of the null distributions by the vertical black lines. The theoretical density of $Q_X$ (see Material and Methods) is shown as a blue curve.

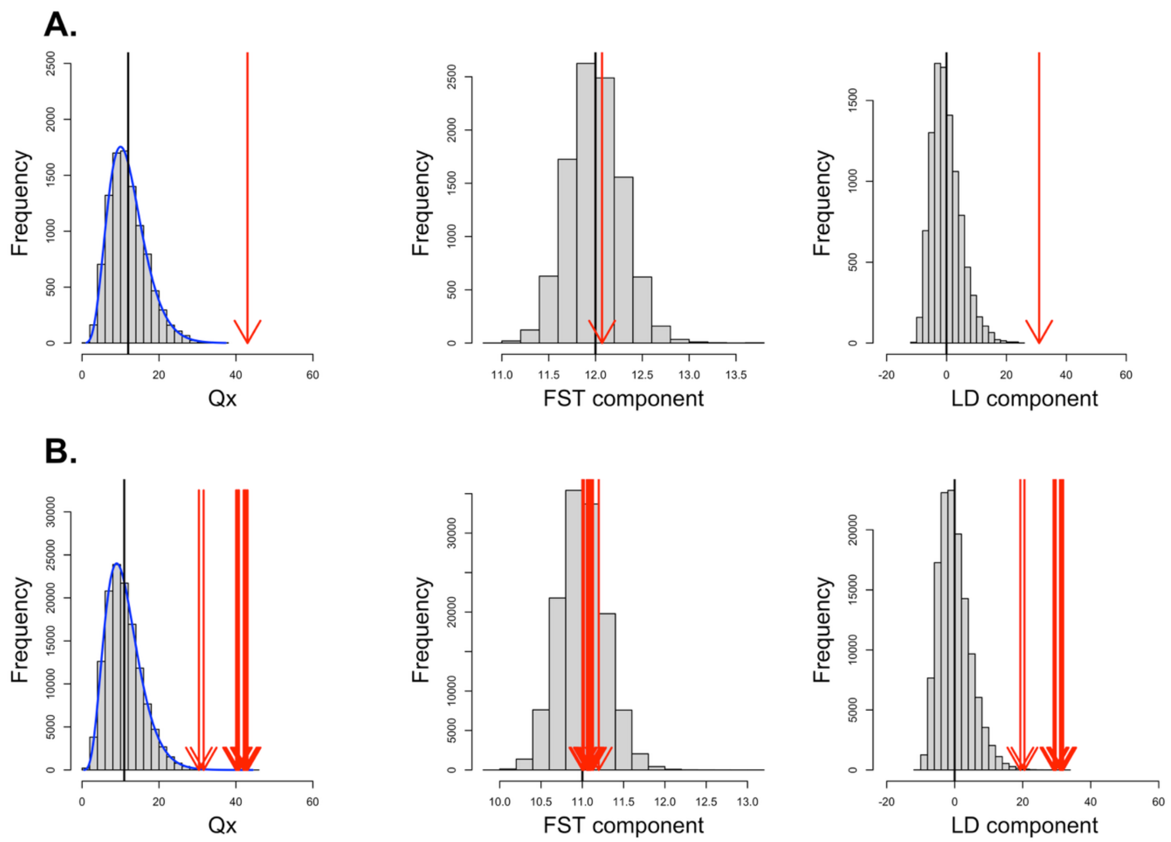


**Supplementary Figure 4: $Q_X$ analysis using 676 EAS COJO SNPs from Yengo et al.** [4]
Histogram of the empirical null distribution of $Q_X$, as well as its $F_{ST}$ and LD component are displayed for A) analysis including the entire set of 13 European populations and B) leave-one-population-out analysis. Null distributions were obtained by genome-wide resampling of SNPs matched to the COJO SNPs by MAF and LD-score. Estimated values are depicted with red arrows, and estimated mean values of the null distributions by the vertical black lines. The theoretical density of $Q_X$ (see Material and Methods) is shown as a blue curve.

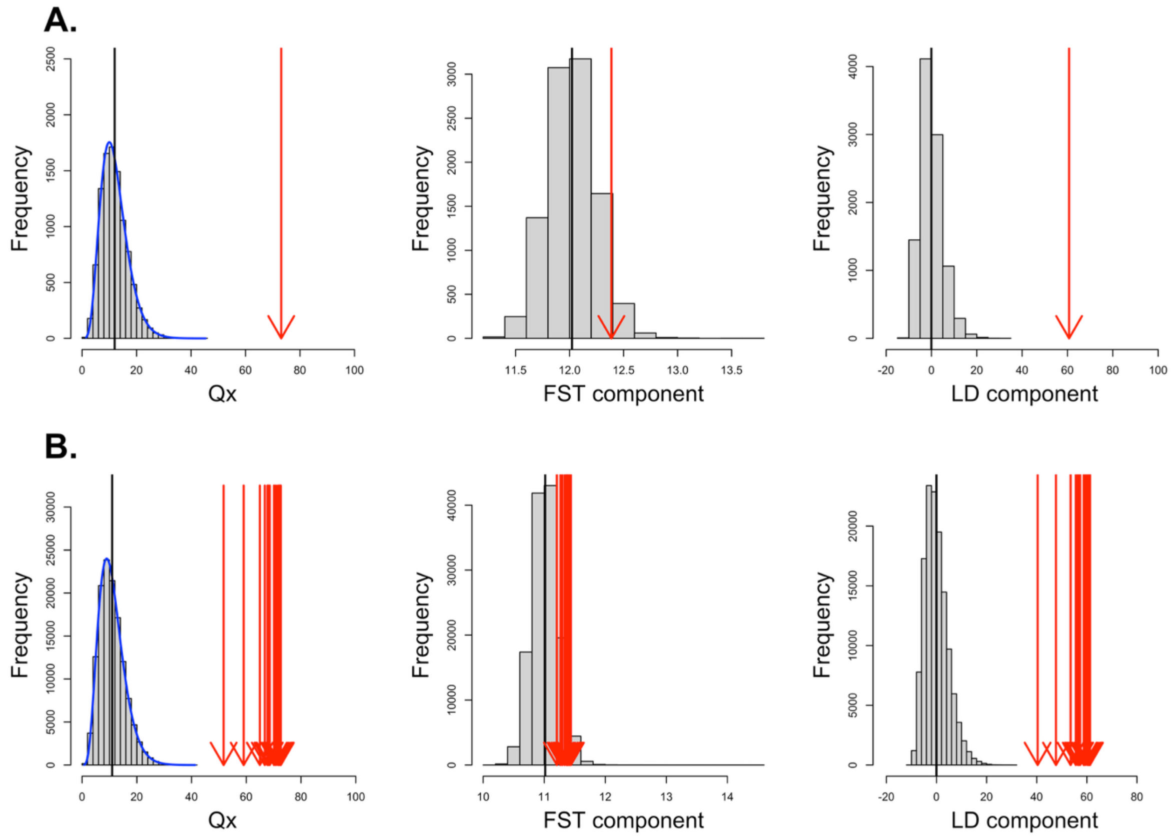


**Supplementary Figure 5: $Q_X$ analysis using 1,100 HIS COJO SNPs from Yengo et al.** [4]
Histogram of the empirical null distribution of $Q_X$, as well as its $F_{ST}$ and LD component are displayed for A) analysis including the entire set of 13 European populations and B) leave-one-population-out analysis. Null distributions were obtained by genome-wide resampling of SNPs matched to the COJO SNPs by MAF and LD-score. Estimated values are depicted with red arrows, and estimated mean values of the null distributions by the vertical black lines. The theoretical density of $Q_X$ (see Material and Methods) is shown as a blue curve.

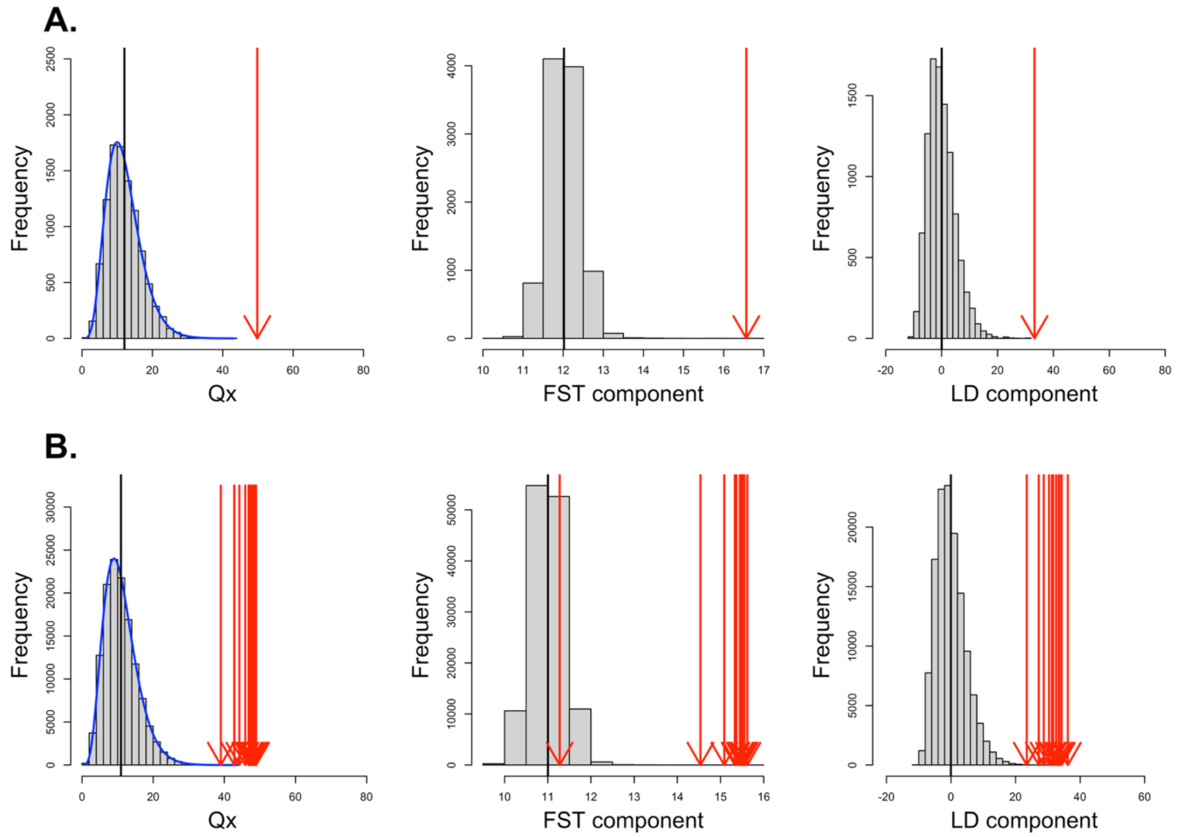


**Supplementary Figure 6: $Q_X$ analysis using 228 EUR COJO SNPs from Howe et al.[10]**
Histogram of the empirical null distribution of $Q_X$, as well as its $F_{ST}$ and LD component are displayed for A) analysis including the entire set of 13 European populations and B) leave-one-population-out analysis. Null distributions were obtained by genome-wide resampling of SNPs matched to the COJO SNPs by MAF and LD-score. Estimated values are depicted with red arrows, and estimated mean values of the null distributions by the vertical black lines. The theoretical density of $Q_X$ (see Material and Methods) is shown as a blue curve.

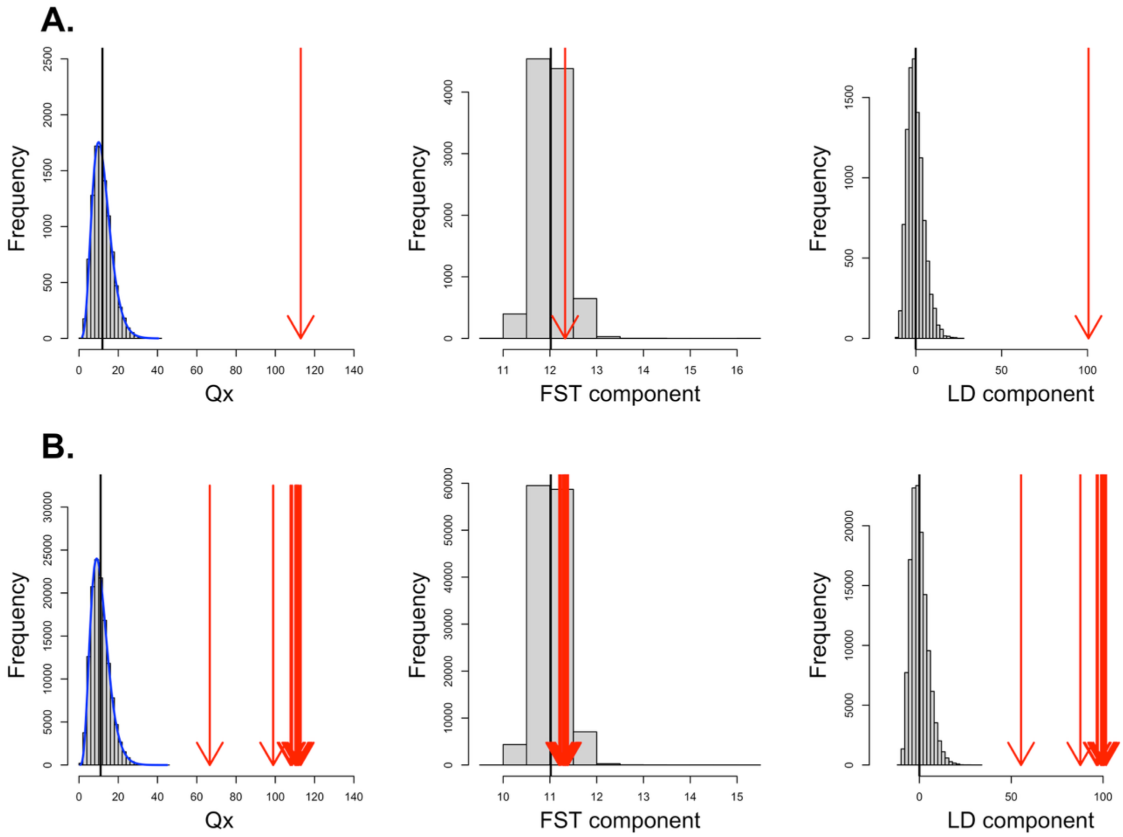


**Supplementary Figure 7: $Q_X$ analysis using 1489 ascertained EUR COJO SNPs from Yengo et al.[4] present in independent LD-blocks[11].** Histogram of the empirical null distribution of $Q_X$, as well as its $F_{ST}$ and LD component are displayed for A) analysis including the entire set of 13 European populations and B) leave-one-population-out analysis. Null distributions were obtained by genome-wide resampling of SNPs matched to the COJO SNPs by MAF and LD-score. Estimated values are depicted with red arrows, and estimated mean values of the null distributions by the vertical black lines. The theoretical density of $Q_X$ (see Material and Methods) is shown as a blue curve.

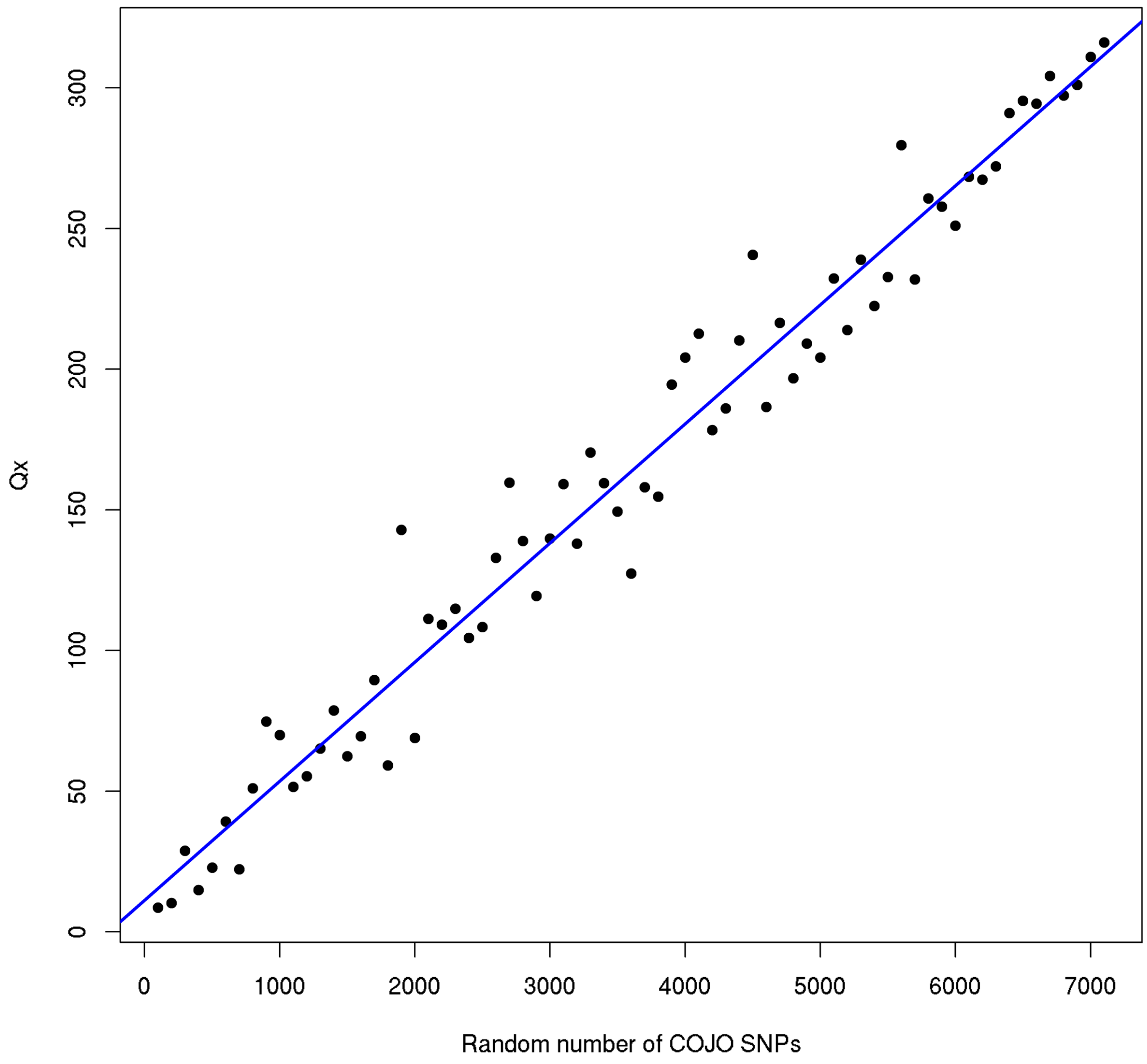


**Supplementary Figure 8: $Q_X$ analysis for different numbers of randomly sampled EUR COJO SNPs from Yengo et al.**[4] COJO SNPs were randomly sampled amongst 7,172 SNPs associated with height in European ancestry GWAS and $Q_X$ analysis included the entire set of 13 European populations. The solid blue line depicts the $Y = 11.25 + 0.042X$ line.

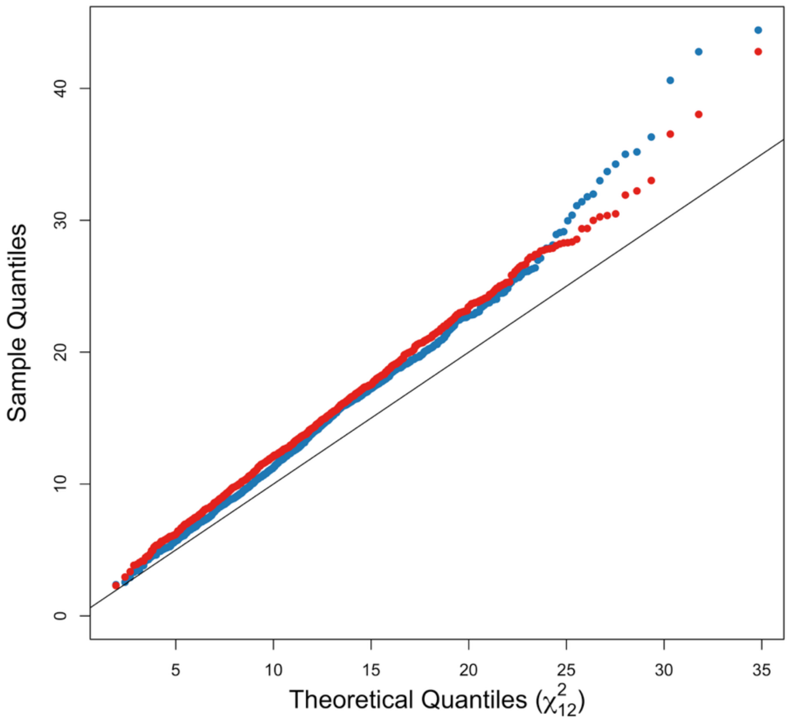


**Supplementary Figure 9: Quantile–quantile (QQ) plots of observed $Q_X$ statistics versus theoretical chi-square quantiles under the null hypothesis of genetic drift, for the assortative mating simulation study (see Materials and Methods).** Results are shown after 10 (blue dots) and 100 (red) generations. The x-axis shows the theoretical quantiles of the chi-square distribution with 12 degrees of freedom, while the y-axis shows the observed $Q_X$ values from simulations (with 1,000 replicates per simulation setting). The solid black line depicts the Y=X line. Upward deviations from this line suggest inflation of the test statistic.

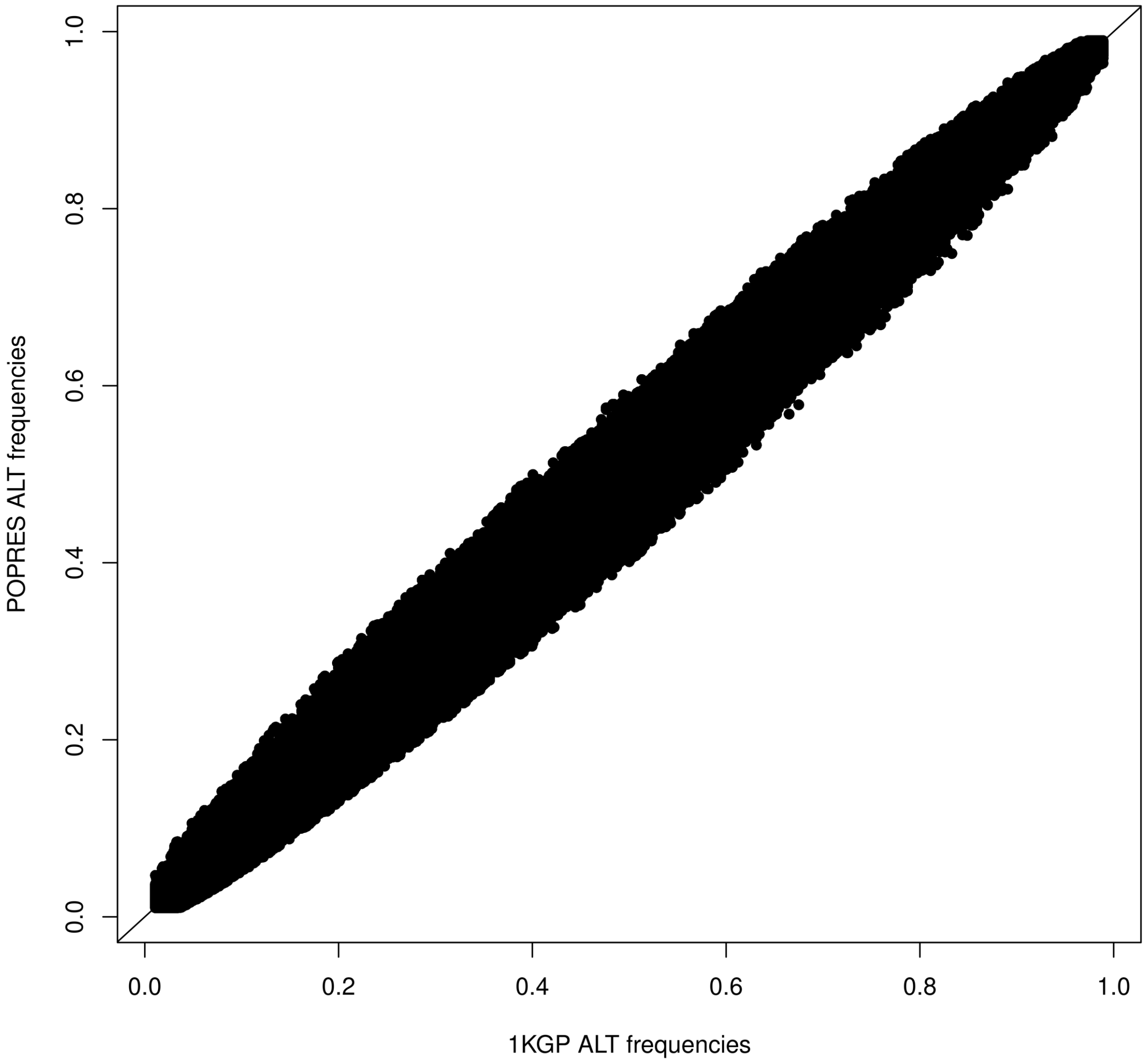


**Supplementary Figure 10: Comparison of alternative allele frequencies of 1KGP and POPRES dataset at 6,253,830 imputed SNPs.** The solid black line depicts the Y=X line.

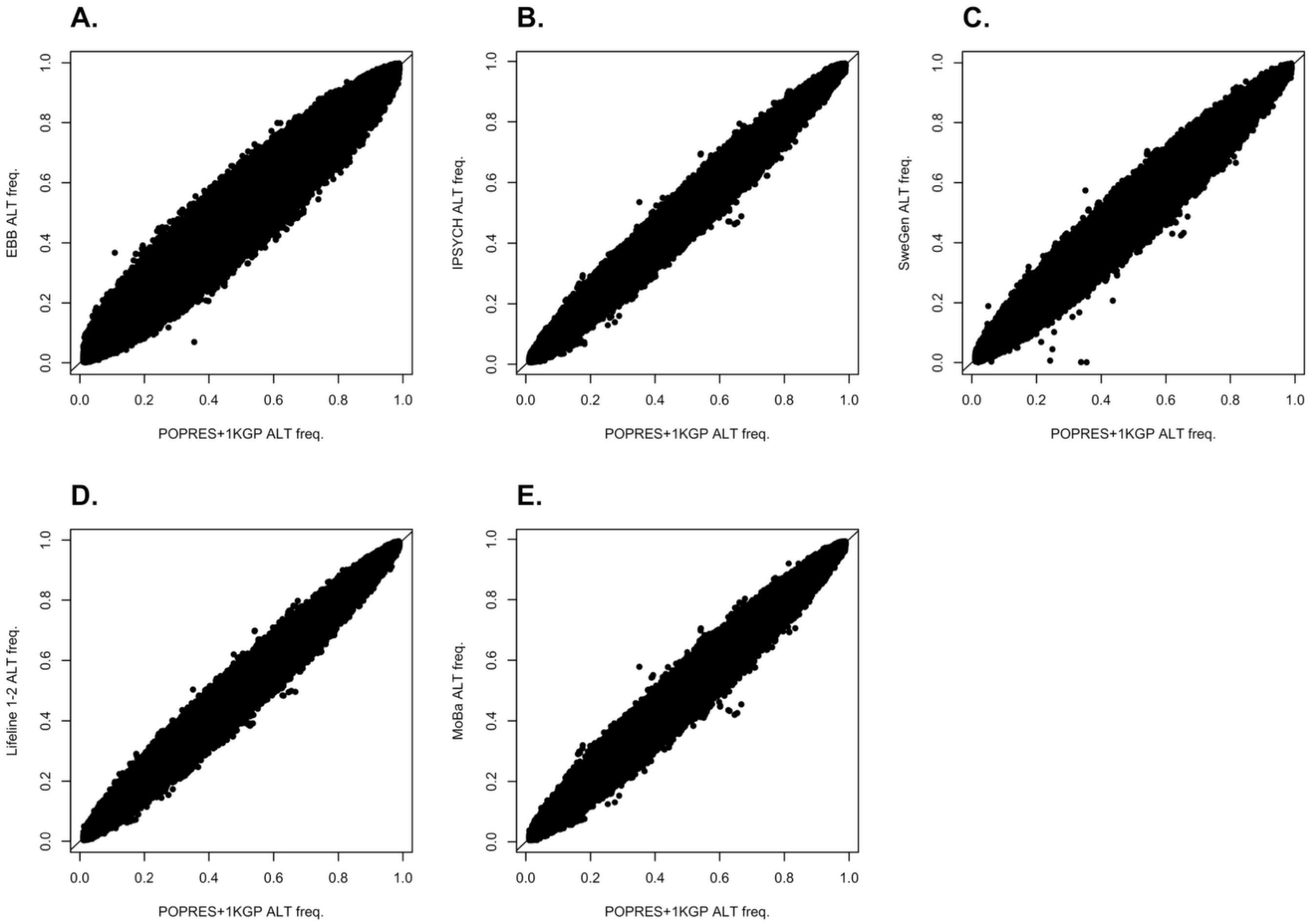


**Supplementary Figure 11: Comparison of allele frequencies between European cohorts.** Alternative allele frequencies at 890,392 SNPs of the POPRES and 1KGP dataset are compared with allele frequencies of A) the Estonian Biobank (EBB), B) IPSYCH, C) SweGen, D) Lifeline 1-2 and E) MoBa cohorts. The solid black line depicts the Y=X line.

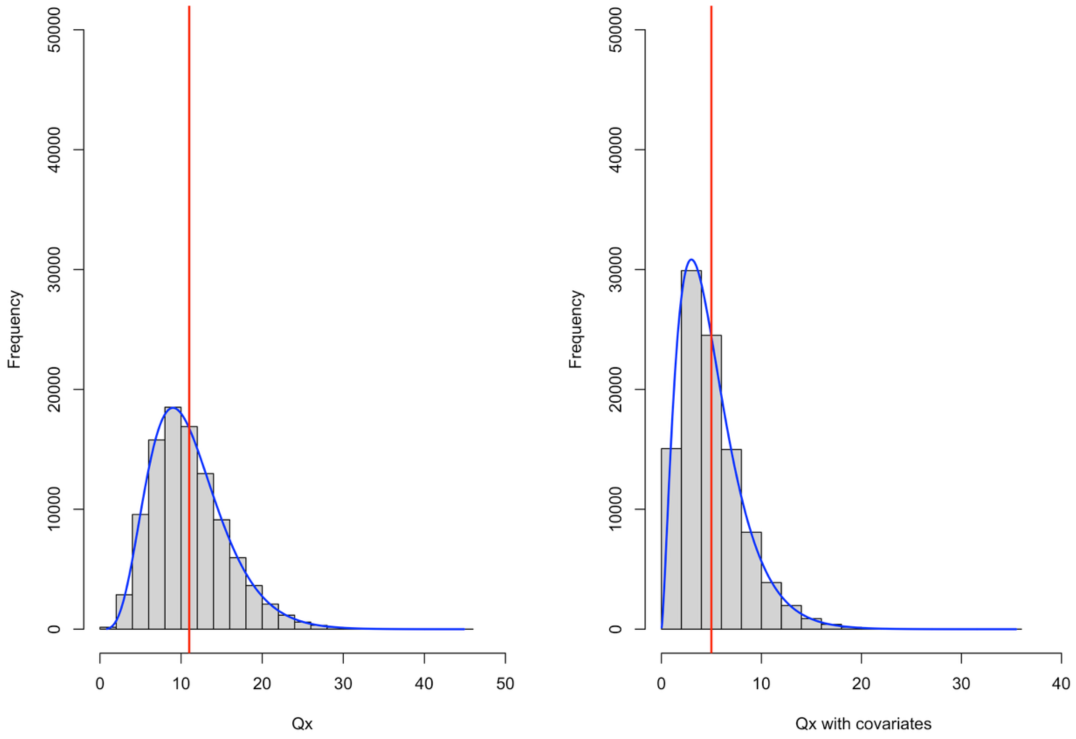


**Supplementary Figure 12: Validation of $Q_X$ corrected for covariates using simulations. We simulated a null dataset using the estimated $F$ matrix on 12 European populations from which we computed $Q_X$ and simulated its null distribution (see Material and Methods) A) without including admixture PCs as covariates and B) when including admixture PCs.** Theoretical mean $Q_X$ values are depicted by the vertical red lines and estimated mean values of the null distributions across 100,000 replicates by the vertical black lines (here, confounded with the red lines). The theoretical density of $Q_X$ (see Material and Methods) is shown as a blue curve.

**Supplementary Table 1: Sample sizes per European country used for allele frequency estimation in the population genetics analysis.**

| Cohort | Sample Size |
|---|---|
| Portugal | 144 |
| Spain (including IBS) | 315 |
| Italy (including TSI) | 339 |
| France | 87 |
| Switzerland | 1058 |
| Netherlands | 64,439 |
| Great Britain | 107 |
| CEU | 183 |
| Denmark | 65,589 |
| Estonia | 49,160 |
| Finland | 105 |
| Sweden | 1000 |
| Norway | 11,093 |

**Supplementary Table 2: Proportion of $Q_X$ statistics explained by 6 admixture PCs for analysis including 12 European populations and leave-one-population-out analysis across five independent set of height associated COJO SNPs.** For each set of populations, the proportion of $Q_X$ explained by the admixture PCs is shown for European (EUR), African (AFR), East Asian (EAS) and Hispanic (HIS) COJO SNPs from Yengo et al.[4], as well as EUR COJO SNPs from Howe et al.[10] within-family GWAS.

| **Population removed** | **SNP set (Number of SNPs)** | | | | |
|---|---|---|---|---|---|
| | Yengo EUR (7,172 SNPs) | Yengo AFR (371 SNPs) | Yengo EAS (676 SNPs) | Yengo HIS (1,100 SNPs) | Howe EUR (228 SNPs) |
| None (12 Populations) | 0.58 | 0.63 | 0.49 | 0.69 | 0.42 |
| Portugal | 0.57 | 0.72 | 0.60 | 0.72 | 0.55 |
| Spain | 0.59 | 0.76 | 0.71 | 0.73 | 0.59 |
| Italy | 0.55 | 0.64 | 0.54 | 062 | 0.48 |
| France | 0.58 | 0.61 | 0.48 | 0.67 | 0.40 |
| Switzerland | 0.59 | 0.64 | 0.6 | 0.69 | 0.46 |
| Great Britain | 0.64 | 0.69 | 0.5 | 0.75 | 0.51 |
| Finland | 0.71 | 0.68 | 0.57 | 0.75 | 0.41 |
| Netherlands | 0.85 | 0.76 | 0.64 | 0.86 | 0.45 |
| Denmark | 0.74 | 0.74 | 0.58 | 0.8 | 0.43 |
| Norway | 0.68 | 0.82 | 0.71 | 0.83 | 0.87 |
| Estonia | 0.66 | 0.64 | 0.55 | 0.71 | 0.44 |
| Sweden | 0.59 | 0.64 | 0.52 | 0.71 | 0.54 |

**Supplementary Table 3: PALM estimates of selection gradients for height within GBR.** Estimated gradient coefficients ± standard errors (p-values) are showed for PALM analysis using respectively 9,079 EUR, 416 AFR and 809 EAS COJO SNPS from Yengo et al.[4] Selection gradient was inferred for the last 2K and 2.7K years before present, assuming a generation time of 29 years. Standard errors were estimated from 5000 block bootstraps.

| **COJO SNP set** | **PALM 2K years bp.** | **PALM 2.7K years bp.** |
|---|---|---|
| Yengo et al. EUR | $0.023 \pm 0.0133$ (0.08) | $0.019 \pm 0.0095$ (0.04) |
| Yengo et al. AFR | $-0.012 \pm 0.0164$ (0.48) | $-0.007 \pm 0.0111$ (0.52) |
| Yengo et al. EAS | $0.009 \pm 0.0180$ (0.62) | $0.007 \pm 0.0127$ (0.57) |
